# Kappa (*κ*): Analysis of Curvature in Biological Image Data using B-splines

**DOI:** 10.1101/852772

**Authors:** Hadrien Mary, Gary J. Brouhard

## Abstract

Curvature is a central morphological feature of tissues, cells, and sub-cellular structures. A challenge for computational biology is to measure the curvature of these structures from biological image data. We present an open-source Fiji plugin for measuring curvature using B-splines. The plugin is named Kappa after the Greek symbol for curvature, *κ*. Kappa is semi-automated: users create an initialization curve by a point-click method, and the initialization curve is fit to the underlying data using an iterative minimization algorithm. We demonstrate Kappa’s applicability on images of cytoskeletal filaments *in vitro*, the cell wall of budding yeast, and whole worms moving in an agar dish. In order to verify the accuracy and precision of Kappa, we created a bank of synthetic images of known curvature using sine waves and golden spirals, which we digitized with different signal-to-noise ratios (SNR), pixel sizes, and point-spread functions (PSF). For synthetic images with characteristics similar to real data, the measured curvatures of those images show a high correlation with the theoretical curvatures. Our fitting algorithms perform better with higher SNR, smaller pixel sizes, and especially PSFs equivalent to super-resolution microscopy data (surprise, surprise). Kappa is freely available under the MIT license for simple integration into Fiji-based workflows. The source code and documentation can be found on GitHub at https://github.com/brouhardlab/Kappa.

## Introduction

Curved structures are found at all scales of biological organization, from whole organisms to organelles to subcellular structures. For example, cell membranes appear in a range of curvatures, including highly-curved clathrin-coated pits (Marsh, 1999; McMahon et al., 2011) and the tubes of the smooth endoplasmic reticulum (ER) (Berridge, 2002; Shibata, Voeltz, et al., 2006). Microtubules and actin filaments are also noticeably curved at the periphery of cells (Brangwynne et al., 2006). Importantly, the curvature of a subcellular structure can be a signal for the recruitment of specific proteins. For example, the Arp2/3 complex binds preferentially to the convex surface of a bent actin filament (Risca et al., 2012), the reticulon protein Dpr1/Yop1 binds to tubular ER membranes (Shibata, Shemesh, et al., 2010), and septins binds to curved plasma membranes (Field, 1996; Pan et al., 2007; Bridges et al., 2016). Our personal interest is in microtubule-associated proteins (MAPs) that bind to curved microtubules, such as TPX2 (Roostalu et al., 2015), Tau (Samsonov et al., 2004), and doublecortin (DCX) (Bechstedt et al., 2014). In order to understand these phenomena, we need to use microscopes to image curved structures, measure their curvature (*κ*), and correlate this curvature with the recruitment of specific proteins. Thus, accurate and precise measurement of curvature is an important requirement for quantitative cell biology and biophysics.

Biological image data are noisy and often crowded, meaning that curved structures within the images may overlap with other features, may vary considerably in intensity along their length, and may change in curvature. Moreover, subcellular structures, such as cell membranes and cytoskeletal filaments, have physical dimensions below the diffraction limit of the light microscope. Images of these structures will be convolved by the optical transfer function (OTF) of the microscopy system. Additionally, modern digital images have pixel sizes on the order of 100 nm, meaning that the OTF is spread over 9–16 pixels for 2D acquisition, which creates a digitization problem (Worring et al., 1993). These conditions can make it difficult to measure the curvature of structures in biological images.

Our goal is to develop software that accurately measures the true curvature of biological structures. Here we provide a robust and easy-to-use Fiji plugin for measuring curvature in one color channel and for correlating this curvature with the intensity in another color channel. We named the plugin Kappa after the Greek symbol for curvature, *κ*. Our plugin uses cubic B-splines, which are parametric piecewise 3rd degree Bézier curves (see Design and Implementation). Bézier curves were developed between 1958 and 1960 by Pierre Bézier and Paul de Casteljau for the French automobile industry (Bézier, 1968). Figure 1A shows a photo of the Citroën 2CV, a car that was redesigned in 1963 using parametric curves. Bézier curves and B-splines are now widely used in computer graphics and computer-aided design (see Fig. 1B), e.g., in the Pen tool in Adobe Illustrator™or in the Bezier Curves tool in Inkscape (Bah, 2011). Because of the ubiquity of B-splines, many fast and numerically stable algorithms exist for plotting B-splines and storing them in efficient data structures (Hoschek et al., 1993). B-splines are locally adjustable and can model complex shapes with a small number of defined points, making them more effective in curve fitting than other parametric curve systems (Rogers, 1977). These properties greatly facilitate the use of B-splines to measure curvature in image data.

**Figure 1.**
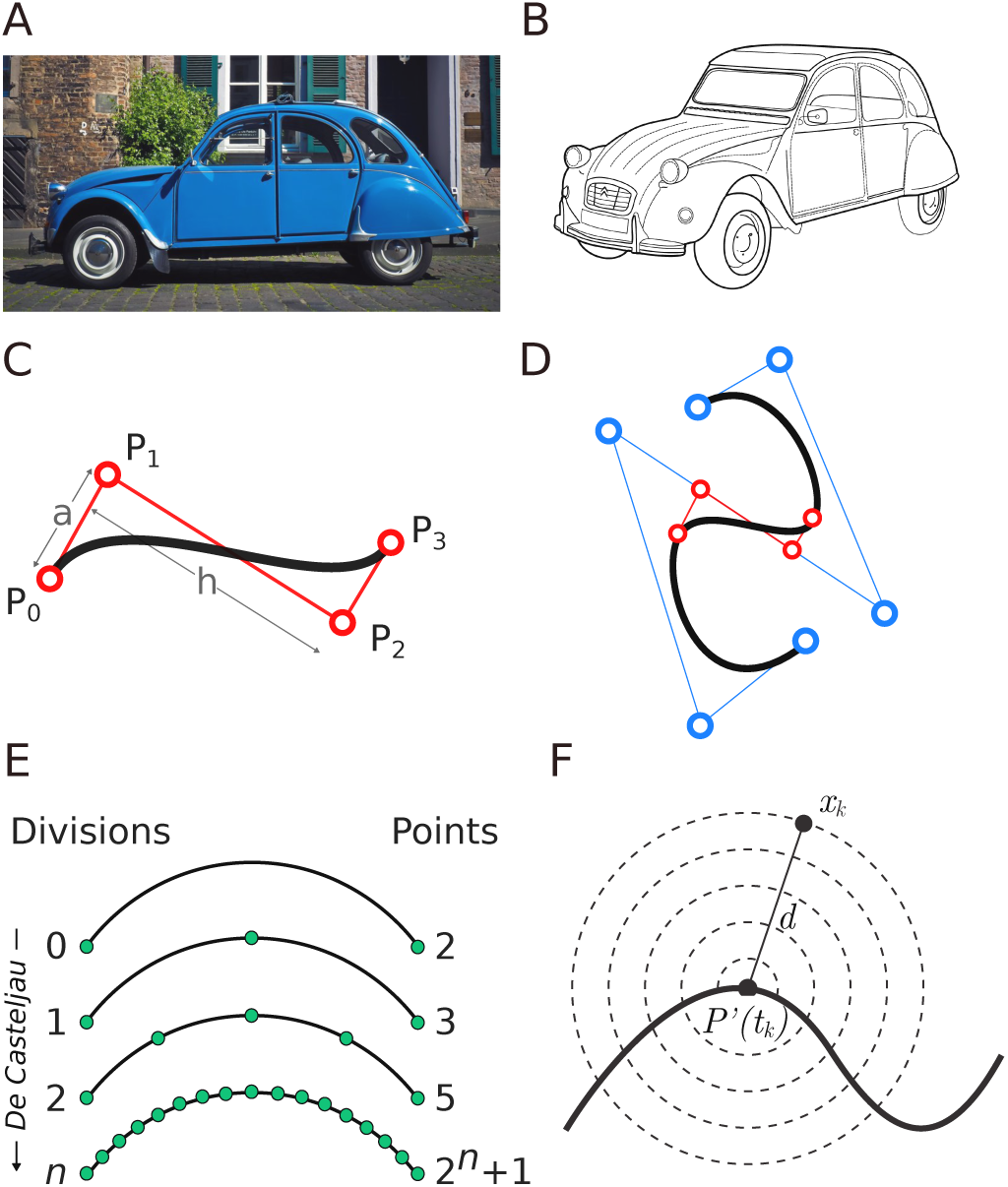
B-splines are easily adapted for curvature analysis. **(A)** Photograph of a blue Citroën 2CV automobile. **(B)** Illustration of a Citroën 2CV generated using B-splines. **(C)** Schematic of a cube Bézier curve (*d* = 3). The curve is shown in black, its 4 control points *P*_*i*_ in red. The line segments *a* and *h* from Eq. 7 are shown in gray. **(D)** Schematic of a B-spline composed of multiple Bézier curves. The blue circles connected by thin blue lines represents the control points drawn by the user. The red control points represent one Bézier curve within the B-spline. **(E)** Diagram of the de Casteljau algorithm that recursively subdivides a Bézier curve *n* times. **(F)** Diagram of iso-values for the point-distance error term (Eq. 8). Data points that are equidistant from a curve point would have the same point-distance error term.

In Kappa, a user draws an initialization curve by clicking along the object of interest. This initialization curve is fit to the underlying data using an iterative minimization algorithm (Plass et al., 1983). Using synthetic images of sine waves and golden spirals, whose curvature is known analytically, we demonstrate that Kappa’s fitting algorithm is accurate and precise. For sine waves, the Pearson correlation coefficient between the true curvature and the fit from Kappa is high (*r* > 0.8) for SNR ≥ 10 dB and across a range of commonly used pixel sizes. An earlier version of this software was used to correlate the binding of Doublecortin to curved microtubules (Bechstedt et al., 2014). The version of Kappa presented here is open source under the MIT license; the source code can be found on GitHub.

https://github.com/brouhardlab/Kappa

## Design and Implementation

### Bézier Curves and B-splines

Bézier curves are parametric curves based on Bernstein polynomials (Davis, 1975). The central concept of Bézier curves is the control point (Fig. 1C). Control points are common to many parametric curve systems. For example, cubic Bézier curves have 4 control points (*P*_*i*_): the first and last control points, *P*_0_ and *P*_3_, are the end points of the curve, while the other points, *P*_1_ and *P*_2_, don’t necessarily lie on it (Fig. 1A). The location of the control points in the *xy*-plane defines the position and shape of the curve.

Most shapes are too complex to define using a single Bézier curve. Complex shapes can be defined using B-splines, which are piecewise sequences of Bézier curves joined at their endpoints. B-spline curves are defined by:

1. The degree *d* of the B-spline, which is also the degree of the underlying Bézier curves.
2. A set of *n* control points *P*_0_, …, *P*_*n*−1_.
3. A knot vector **U** = [*u*_0_,…, *u*_*n*+*d*_] defining the parameter values for the individual Bézier curves that comprise the B-spline.
4. A set of B-spline basis functions, 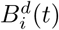.

Given these definitions, the corresponding B-spline curve **B** is given by:

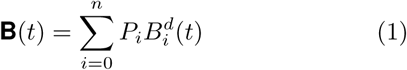

The basis functions 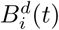 are defined recursively by the de Boor formula:

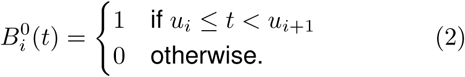

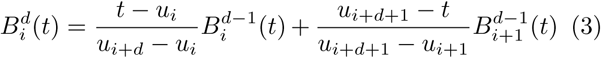

where 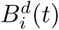) is the *i*^*th*^ basis function of degree *d*. Kappa uses cubic B-splines (*d* = 3), which are the most common type, and thus these basis functions are 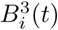. The Bézier curves within B-splines are joined automatically with **C**^*d*−1^ continuity (Figure 1B); thus, curve junctions for cubic Bézier curves have **C**^2^ continuity, meaning that the first and second parametric derivatives of the two curve sections are identical at the junction, ensuring smoothness.

In Kappa, the user draws control points on an image by clicking along the length of a curved region of interest. These clicks will produce an “initialization curve.” The initialization curve can be adjusted by clicking and dragging the control points. This straightforward manipulation of B-spline curves is a major reason for their popularity and an advantage over piecewise polynomial curves. For sparse images, the initialization B-spline doesn’t need to follow the curved region of interest on the image closely since it can be fit later to the underlying data (see below). For dense images, greater care is encouraged.

### Curvature Measurement

The curvature *κ* at a point on the line can be intuitively defined as the inverse of the radius of curvature:

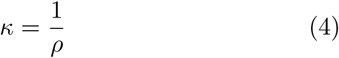

where *ρ* is the radius of the circular arc *C* whose tangent vector is the same as the curve’s tangent vector at that point. For a straight line, *ρ* = ∞ and *κ* = 0, whereas a circle with *r* = 2 *μm* has *κ* = 0.5 *μm*^−1^. Thus, a simplistic way to measure curvature is to draw circles on an image and measure their radii. This method works best when the curvature *κ* is constant. For lines with variable curvature, approximating the circular arc *C* for a specific point on the curved line can be difficult. So the curvature *κ* can also be defined as the speed at which the unit tangent vector 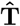 rotates for a given segment on the curve *s* (Pressley, 2010):

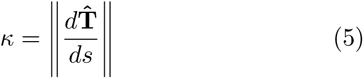

Then for a parametric curve in the *xy*-plane, the absolute curvature, *κ*, is defined as:

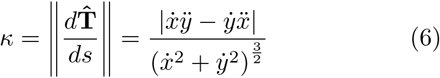

Evaluating the curvature of a Bézier curve from Eq. 6 as a function of *t* is possible but mathematically complex (Sederberg, 2012). But the advantage of Bézier curves is that Eq. 6 is mathematically simple when evaluated at an endpoint *t*_0_ of a Bézier curve (Sederberg, 2012):

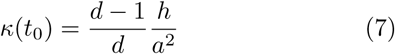

where *d* is the degree of the curve, *a* is the length of the line segment between the first two control points (*P*_0_ and *P*_1_) and *h* is the perpendicular distance between the third control (*P*_2_) point and the secant passing through the first two control points (see Fig. 1C for illustration). For B-splines, we can easily calculate *κ* at curve junctions, where two Bézier curves are joined at their endpoints.

Therefore, a fast and easy way to measure the curvature along the length of the B-spline curve is to extract its Bézier curves using the Böhm algorithm (Sederberg, 2012; Böhm, 1981) and calculate the curvature at the endpoints of these curves using Eq. 7. Furthermore, we can take advantage of the well-known de Casteljau algorithm (Boor, 1978; Böhm, 1981), in which any Bézier curve can be divided in two. This procedure creates additional endpoints at the new curve junctions, at which the curvature can be measured using Eq. 7. By recursing this subdivision event *n* times, we obtain 2^*n*^ + 1 end-points, which are used to provide a fine-grained measurement of curvature along the entire B-spline curve (see Fig. 1C). As mentioned above, cubic B-splines are well-suited to this type of analysis because they are **C**^2^ continuous. In Kappa’s interface, an information panel plots the fine-grained curvature as a function of length along the B-spline and reports the mean curvature along the B-spline curve as well as the curvature at any point on the curve selected by the cursor.

### Curve Fitting

Manually-adjusted B-splines may be prone to selection bias when clicked out and adjusted. In order to minimize this bias and obtain consistent and reproducible curvature measurements, the initialization B-spline curve can be fit to the data in the image. This step is not mandatory and users can skip it when making casual measurements.

In brief, the positions and intensity values of pixels that represents the curved feature are used to fit a B-spline curve. The pixel selection procedure is done by specifying two parameters:

1. The width around the initialization curve to include, which defines the fitting area.
2. A brightness threshold for data to include. The threshold can be toggled to choose brighter pixels against a dark background or darker pixels against a white background, depending on the type of data. In both cases, the fitting procedure is weighted by pixel brightness above or below the threshold (see below).

The fitting area is traced on the image in light blue (see Fig. 2). Pixels that will be included in the fitting procedure are colored pink in the image.

**Figure 2.**
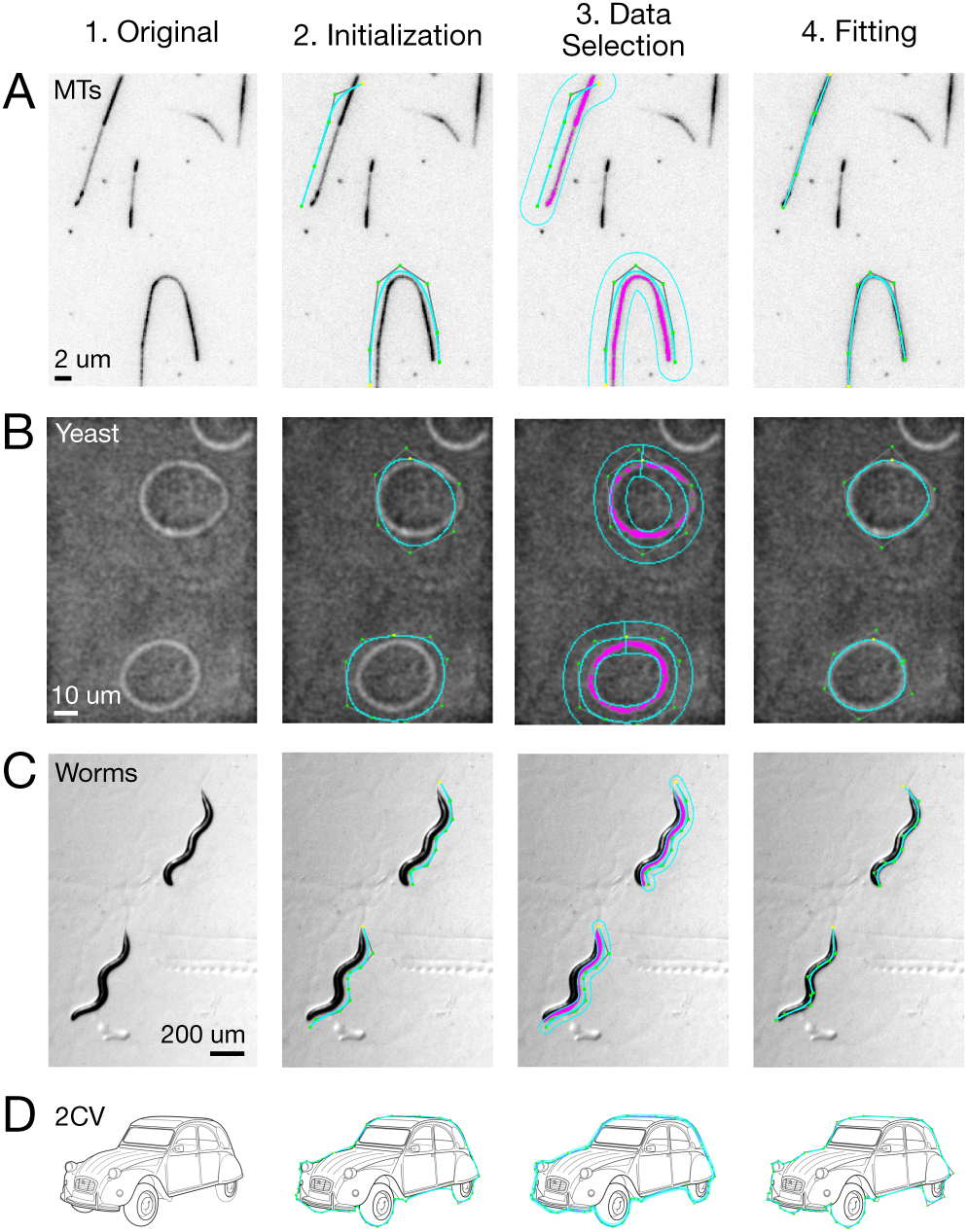
Kappa can be used on a wide range of images. Column (1) shows the original images. In column (2), the blue lines are initilization curves drawn by a user. Column (3) shows the selection of data for fitting; the fitting area is outlined in blue and the selected pixels are pink. In (4), the post-fitting B-spline is shown in blue. **(A)** Images of curving rhodamine-labeled microtubules attached to a cover glass surface. **(B)** Images of budding yeast, *S. cerevisiae*. **(C)** Images of *C. elegans* on an agar plate. **(D)** Outline of a Citroën 2CV.

To fit the curve to the pink pixels, we employ an iterative minimization algorithm based on least-squares methods (Hoschek et al., 1993). This approach is widely used in surface fitting and can be applied to any parametric curve (Grossman, 1971). When applied specifically to B-splines, several unique implementations have been published. We implemented two of these. The first method uses a spatial error term, described below, that is iteratively reduced (Hoschek, 1988) to bring the B-spline closer to the data; originally called “intrinsic parametrization for approximation”, this method is nowadays known as “point distance minimization” (PDM). A later approach known as “squared distance minimization” (SDM) uses a more complex, curvature-dependent error term, which converges more rapidly than PDM (Wang et al., 2006).

We observed very similar performance for PDM and SDM in tests of synthetic image data, described below. Both PDM and SDM are available as options in the software. Other approaches, such as “tangent distance minimization” (TDM) (Blake et al., 2012), could be part of future development.

PDM involves updating control points of an initial B-spline curve so as to minimize the Euclidean distance between the current fitting curve and each data point *X*_*k*_. In other words, it minimizes the local error term *e*_*P,k*_:

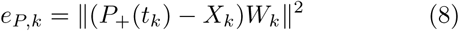

where *P*_+_(*t*_*k*_) is the fitting curve with updated control points, *X*_*k*_ are the positions of pixels and *W*_*k*_ are the weights for each *X*_*k*_, measured as the square root of the *X*_*k*_ pixel intensities. Figure 1D illustrates iso-value curves for the point distance error term.

While the concept of minimizing a Euclidean distance is intuitive, we do not know which data point *X*_*k*_ should be minimized against which curve point *P*(*t*_*k*_) on the B-spline. The optimal curve point *P*(*t*_*k*_) is called the *footpoint* of *X*_*k*_ on the curve *P*(*t*). The optimal footpoints are not known *a priori*, however. We adopted the common method of choosing the initial B-spline parameters *t*_*k*_ such that *P*(*t*_*k*_) is the closest point from the current fitting curve to the data point *X*_*k*_. In other words, we start with the smallest Euclidean distance between data points and curve points.

After mapping each data point *X*_*k*_ to its nearest foot-point *P*(*t*_*k*_), the average global error *E*_*k*_ is computed:

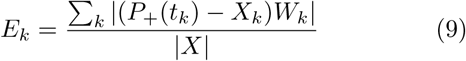

where |*X*| is the number of data points considered. We apply an iterative method until convergence to a local minima by interleaving footpoint computation with the minimization of the error term *E*_*k*_, where each iteration updates the control points *P*_+_. This method produces a locally-optimal fitted curve in polynomial time.

### Control Point Removal

Besides minimizing the error term, our algorithm also checks for over-fitting, one of the major pitfalls of parametric curve fitting. Over-fitting occurs when the fitted curve has an excessive number of parameters that exaggerate small fluctuations in the underlying data. For example, Bicek et al. observed over-fitting when they measured the curvature of cytoskeletal filaments such as microtubules (Bicek et al., 2007). More specifically, they observed that shape fitting using sums of cosine waves or higher order polynomials performed poorly when the initialization curve was overly complex. In the case of B-splines, over-fitting occurs when a curve has too many control points: if too many points are used to define the initialization curve, the fit may produce large variations in curvature within over-fitted regions. Since the optimal number of control points is not known *a priori*, we need a method to reduce the complexity of the B-spline curve, when necessary, during fitting.

Several methods have been used to reduce the number of control points. These methods include algorithms that penalize the fit against higher variations in curvature (Eilers et al., 1996) and post-fitting adjustment algorithms where the number and distribution of control points is adjusted after the curve is fit (Hoschek, 1987; Yang et al., 2004). We chose an approach inspired by the latter set of algorithms. By assuming that our initial fit broadly matches the shape of the underlying curve, we make small adjustments to specific regions that are potentially over-fit. To identify these regions, we use a simple heuristic, namely that areas where the control points are densely spaced are prone to over-fitting. We iteratively identify and remove these “dense” control points while evaluating if the global and local errors of the fit remain within ranges defined by the user.

To locate the “densest” point, we identify the index of the control point that has the smallest Euclidean distance to its nearest neighbor. If this is the *i*^*th*^ control point, we replace the *i*^*th*^ and (*i* + 1)^*th*^ control point with a new control point whose coordinates bisect the line segment formed between the two original points. In other words, if our current fitted curve has a set of control points *P*_*j*_:

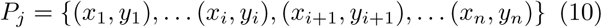

then we replace it with an adjusted curve with control points *P*_*j*+1_:

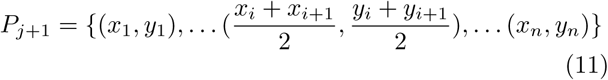

If the “densest” control point is the last control point, we instead do this with the (*i* − 1)^*th*^ and *i*^*th*^ points.

We perform our curve-fitting algorithm on the reduced curve and determine if it still fits the data acceptably. To determine acceptability, we set two default criteria: (1) the global error cannot increase by more than a user-defined value *ε*_*G*_ (Eq. 12), and (2) the maximum local error for any piece of the B-spline cannot increase by more than a user-defined value *ε*_*L*_ (Eq. 13).

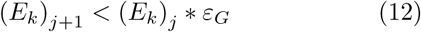

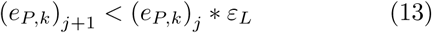

Where 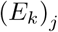 and 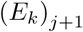 are the global errors before and after curve adjustment. The same definitions apply for the maximum local error, *e*_*P,k*_. We chose relative thresholds as opposed to absolute thresholds so control point adjustment can be applied to a range of images without modification. Default values for those thresholds have been defined empirically as *ε*_*G*_ = *ε*_*L*_ = 1.05. If needed, error thresholds can be adjusted directly in Kappa’s interface. E.g., images with highly curved structures may benefit from a smaller error threshold in order to prevent too much smoothing of the curve.

### Correlating Image Intensity with Curvature

In many cases, the question of interest is whether a particular protein is recruited by or enriched on regions of curvature, and more generally whether a protein’s concentration can be correlated with curvature. By imaging the curved substrate in one color channel (e.g., a cell membrane or a cytoskeletal filament) and the protein of interest in a second color channel (e.g., DCX or a protein that binds to membranes), the user can determine whether fluorescence intensity correlates with curvature. For example, in an previous study our lab used Kappa to show that Doublecortin (DCX), a microtubule-associated protein, binds with higher affinity to curved microtubule lattices (Bechstedt et al., 2014). Kappa shows the intensities for the pixels directly below the selected curve in the user interface, allowing the user to quickly check for a correlation. Data can also be exported for more detailed analysis outside of Kappa (see next section).

### Import / Export

Since Kappa is a plugin for Fiji (Schindelin et al., 2012), it can open a wide range of image formats, including TIF, JPG, PNG, AVI and ND2. The interface allows a user to play through a multi-image movie file and zoom into regions of interest as required. For example, the program can open 16-bit, 3-color TIFs and display each color channel independently. Fig. S1 shows a screenshot of the user interface.

All data generated by Kappa (curves, curvature values, and intensities) can be exported in a comma-separated values file (CSV) for later analysis in almost any programming language or software, such as Python, Julia, Matlab, or Excel. The control points of the B-splines can also be exported in a custom file format (`.kapp`). Note that we also added a feature to import curves from the ImageJ Roi Manager. Therefore, depending on the application, it’s possible to pre-compute curves positions, import them in Roi Manager, and then use them in Kappa (with or without applying the fitting algorithm). Python-based examples of a writer and a reader function for `.kapp` files are available at https://github.com/brouhardlab/Kappa.

## Validation

### Kappa works for a wide range of image types

To demonstrate the applicability of our software, we measured the curvature of structures in several types of images. Our personal motivation for Kappa’s development was to measure the curvature of cytoskeletal filaments such as microtubules (Bechstedt et al., 2014). Figure 2A shows a fluorescence microscopy image of fluorescently-labeled microtubules attached to a cover glass surface that curve naturally during the process of surface attachment (Bechstedt et al., 2014). Column 1 shows the original image. Column 2 shows initialization curves for 2 microtubules created by the point-click method; notice that these initialization curves do not match the microtubules closely. Column 3 shows the area around the initialization curves that will be included in the fitting procedure (blue outlines) and the data that will be included in the fits based on a user-defined threshold (pink). Finally, column 4 shows the curves after the fitting procedure is complete.

Another application of our software is to measure the curvature of cell walls and membranes such as the plasma membrane. As an example, Figure 2B shows an bright-field microscopy image of budding yeast cells, with the columns corresponding to the sequence described above. Moving up in size, Figure 2C shows a dissection microscope image of *C. elegans* moving on an agar plate. Kappa can measure curvature along the length of the worm, which can be used for behavioural or mobility studies. Lastly, Figure 2D shows the outline curves of a Citroën 2CV. In principle, any clearly labeled object contour can be analyzed within an image.

### Generating Synthetic Image Data

In order to assess Kappa’s accuracy and precision, we used Python to create two different datasets that contain images of objects of known curvature, namely sine waves and golden spirals. Sine waves are an excellent test case because they are approximately linear in their middles but highly curved around their peaks, a pattern that replicates the curvature variability seen, e.g., in our microtubule data sets.

Sine waves are defined by a period *T* and an amplitude *A* according to:

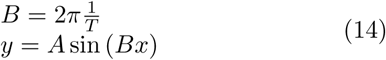

Using Eq. 5, we show the curvature of a sine wave with an amplitude *A* and a period *T* is defined as:

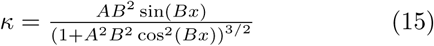

For the sine wave dataset, sine waves (Eq. 14, see Fig. 3A, column 1) were discretized into an array (Fig. 3A, column 2); the array was imagined as having physical dimensions corresponding to 10 nm per array element, and the sine wave had a corresponding period of *T* = 15 *μm* and an amplitude of *A* = 6 *μm*. The discrete elements in the sine wave array were convolved with a 2D Gaussian function, typically using *σ* = 300 nm, corresponding to *λ* = 530 nm light (e.g., the emission peak of GFP, see Fig. 3A, column 3). Noise was added to the image using an additive Gaussian white noise model where the signal-to-noise ratio can be specified (Fig. 3A, column 4). Finally, the high-density array was binned into larger “pixels” corresponding to common pixel sizes found in light microscopy images. The images in Figure 3A shows the generation of a sine wave with a SNR of 20 dB, a pixel size of 160 nm/pixel, and *σ* = 300 nm. As described below, our software measured their peak curvatures and mean curvatures accurately and precisely.

**Figure 3.**
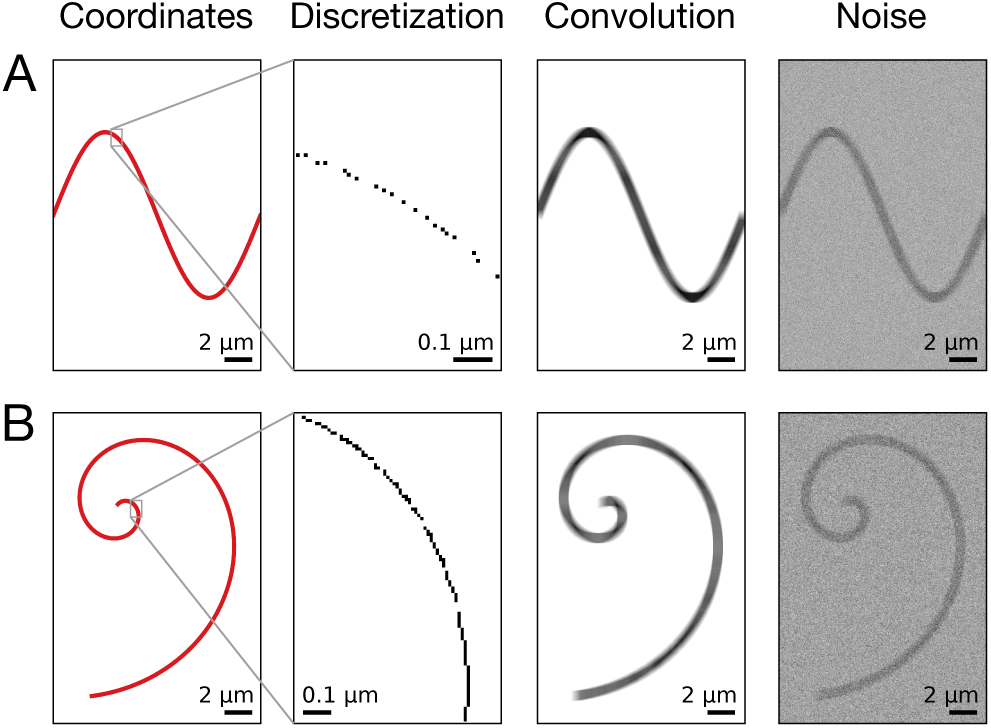
Synthetic image data was generated for sine waves and golden spirals. The steps shown are (1) the generation of coordinate positions from analytical equations, (2) the discretization of the coordinates into pixels of defined size, (3) the convolution of the discrete data with a point-spread function, and (4) the addition of noise. **(A)** Synthetic image of a sine wave, which replicates the curvature found in our images of microtubules. This example has an SNR of 20 dB, a pixel size of 160 nm/pixel, and a PSF of *σ* = 300 nm. **(B)** Synthetic image of a golden spiral with the same values as the sine wave in (A).

Golden spirals are a more complex test case. They are found in the shells of snails and nautilus, the cochlea of the inner ear, hurricanes, and some spiral arms of the Milky Way galaxy. A golden spiral is a special type of logarithmic spiral. The polar equation of a logarithmic spiral is defined as:

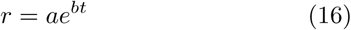

where *t* is the tangential angle of the spiral (it can be thought of as the *x*-axis of the spiral), *a* is the initial radius of the spiral and *b* is the growth factor. The polar form of the logarithmic spiral can be represented in Cartesian coordinates as:

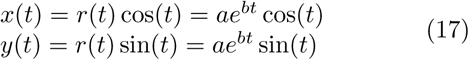

where *x* and *y* are the *xy*-plane coordinates.

In the case of the golden spiral, the growth factor *b* is related to *φ*, the golden ratio,

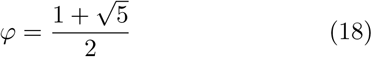

and *b* is defined as

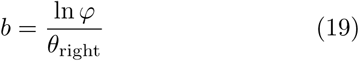

where *θ*_right_ is a right angle (*π/*2 or −*π/*2).

Finally, the curvature *κ* of a logarithmic spiral is given by:

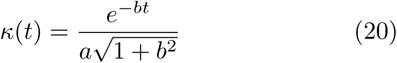

As with the sine wave dataset, golden spiral curves (Fig. 3B, column 1) were discretized with an original pixel size of 10 nm/pixel (Fig. 3B, column 2) and convolved with PSF (Fig. 3B, column 3). As above, we added noise (Fig. 3B, column 4) and pixelated the high-density array. The images in Figure 3B show the generation of a golden spiral with a SNR of 20 dB, a pixel size of 160 nm/pixel, and *σ* = 300 nm. The spirals proved to be a puzzle for our software; our results indicate that the fitting algorithm struggles with their innermost region of rapidly changing curvature.

Jupyter notebooks (Thomas et al., 2016) are available at https://github.com/brouhardlab/Kappa/tree/master/Analysis in order to reproduce both the sine wave and golden spiral datasets. Notebooks to reproduce fitting and curvatures measurements (discussed below) are available in the same repository.

### Kappa is accurate and precise

Using the synthetic images described above, we tested the accuracy and precision of our fitting algorithm. To simulate actual measurement conditions, in which different users will create different initialization curves, we used a set of initialization curves as a starting point for the fitting algorithm (30 curves for the sine dataset and 10 curves for the spiral dataset). These initialization curves varied in control point number and only loosely fit the underlying curve. We analyzed our fitting error three ways: (1) using Pearson’s correlation coefficient *r*, (2) the error in the mean curvature, and (3) the error in the peak curvature for each image type. We determined the error of our curvature measurements as a function of SNR, pixel size, and the position of the initialization curve. In all cases, we observed a very high correlation between the theoretical and measured curvatures.

In order to determine Pearson’s *r*, we need to align our measured curvatures with the theoretical values. For the sine wave dataset, the process is simple and the curvature errors were calculated for each *x* position by taking the absolute difference between the theoretical curvature value from Eq. 15 and the measured curvature value returned by Kappa. For the spiral dataset, the theoretical curvature value can’t be expressed in term of (*x, y*) positions but is expressed from the tangential angle *t* (Eq. 20). We generated theoretical curvature values and (*x, y*) positions from a high-resolution golden spiral curve with a step size of 1 nm. We then aligned the (*x, y*) positions returned by Kappa to the high-resolution (*x, y*) spiral positions by minimizing their Euclidean distances. Once aligned, we take the absolute difference between the theoretical curvature values and the measured curvature values.

#### Variable SNR

To determine if our minimization algorithm is robust against noise, we varied the signal-to-noise ratio (SNR) in the sine wave (Figure 4A) and the golden spiral datasets (Figure 4B) from 1 to 25 dB with a fixed pixel size of 0.16 *μ*m/pixel.

**Figure 4.**
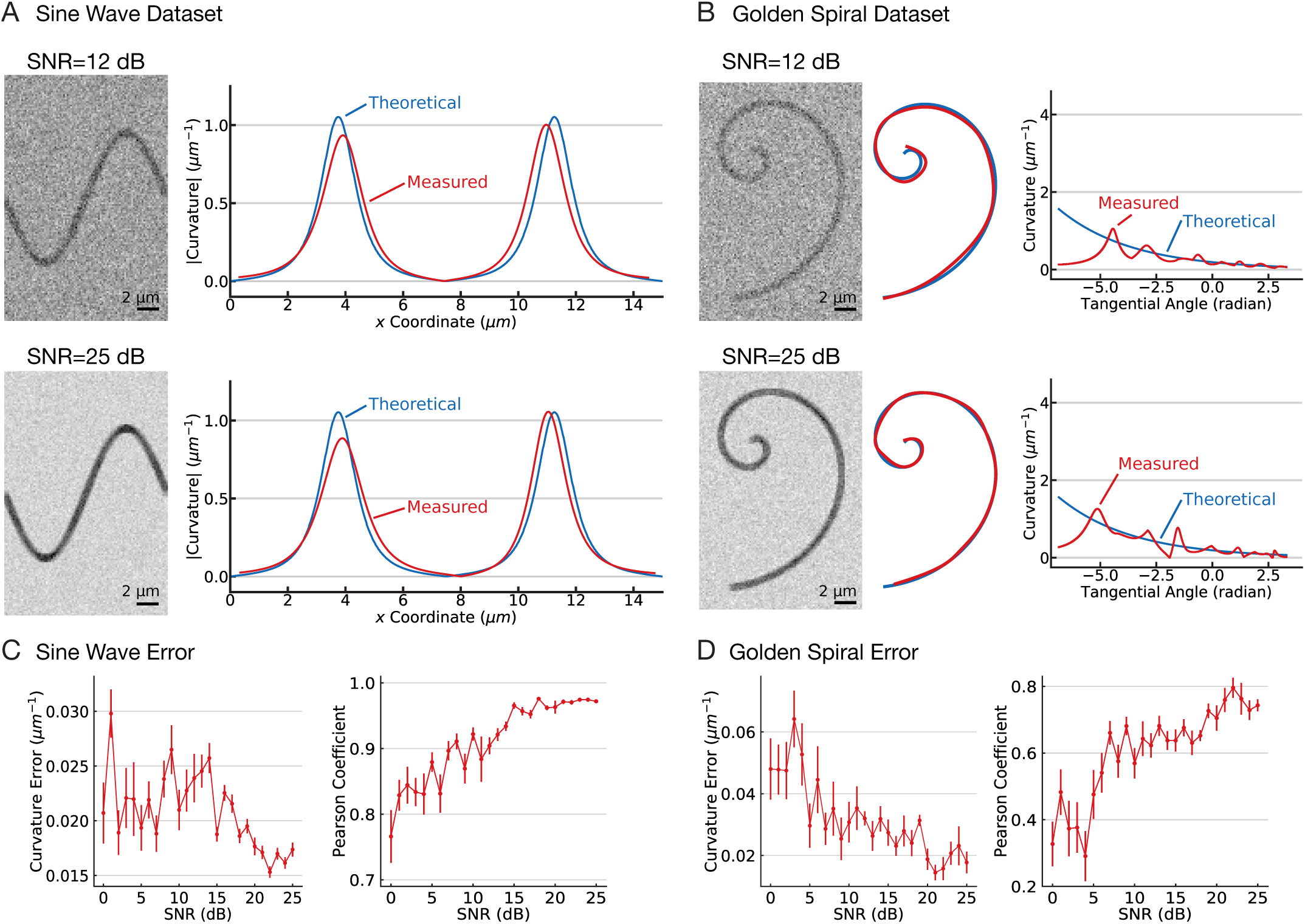
Fitting errors are reduced with increasing SNR. **(A)** Synthetic images of sine waves (*left*) at two SNRs (*labeled*). Plot of sine wave curvature against the *x* coordinate of the curve. The theoretical values are shown in blue and the measured values from a single measurent are shown in red. **(B)** Synthetic images of golden spirals (*left*) at two SNRs (*labeled*). Plot of golden spiral curvature against the tangent angle of the curve (equivalent to a curve length coordinate). The theoretical values are shown in blue and the measured values from a single measurent are shown in red. **(C)** Error in *n* = 30 independent measurements of sine waves. (*Left*) Plot of the error in the mean curvature as a function of SNR. Error bars are the SEM. (*Right*) Plot of the Pearson correlation coefficient as a function of SNR. Error bars are the SEM. **(D)** Error in *n* = 30 independent measurements of golden spirals. (*Left*) Plot of the error in the mean curvature as a function of SNR. Error bars are the SEM. (*Right*) Plot of the Pearson correlation coefficient as a function of SNR. Error bars are the SEM.

Figure 4A,B shows a plot of one such measurement of the sine wave dataset; in this example (and many others), the shape of the measured curvature closely matches the theoretical values with small shifts. While there is general agreement in shape and scale between the two distributions, there is an *x*-offset between their respective peaks. We attribute this offset primarily to the convolution of the data with a PSF, which obscures the true position of the curve. In other words, the shift between our measured distribution and the theoretical distribution is likely systemic and not procedural.

In order to measure the correlation between the theoretical and measured curvatures values, we determined the Pearson correlation coefficient *r* for each fit for the sine wave dataset. Not surprisingly, *r* increases with higher SNR. The correlation peaks at *r* ≃ 0.95 and remains near that point for all SNR >15 dB (Figure 4D). For SNR < 10 dB, the *r* value dropped to a correlation of *r* ≃ 0.75. We conclude that *r* is sufficiently high to make useful measurements across all SNRs.

Pearson’s *r* is sensitive to the *x*-offset discussed above, which exaggerates differences between the true and measured values. In many cases, the precise position of the measured curve is less important than, e.g., the mean curvature of the structure being measured. As such, we determined the error in the mean curvature returned by Kappa for the sine wave dataset. As excepted, we observed a decrease of the mean curvature error as the SNR increases (Figure 4C). Relative to a true mean curvature of *κ* = 0.248 *μm*^−1^, the curvature error at low SNR (< 5 dB) is around 0.022 *μm*^−1^ and decreases to 0.017 *μm*^−1^ for high SNR values (> 20 dB).

Correlation coefficients and mean curvature errors are useful, but they include both straight and curved regions of the data. In analyzing curved structures, it is often the curved areas that are of interest rather than the straight ones. E.g., for the sine wave dataset, points of particular interest are the peaks. We measured a peak curvature of 0.98 ± 0.15 *μm*^−1^ at SNR of 12 dB, which compares favorably to the theoretical curvature of 1.05 *μm*^−1^ at *x* = *T/*4. Thus, we conclude that Kappa accurately and precisely measures both the peak curvature and mean curvature of the sine wave dataset, particularly at high SNR.

For the golden spiral dataset, the curvature changes rapidly near the center, because the curvature depends on the first and second derivative of the shape. We found that Kappa struggled to return smooth curvature values in this region (Figure 4B); instead, the output included a series of peaks in curvature that decayed into a smooth curve. As a consequence, we observe overall higher errors for the golden spiral dataset. The correlation peaks at *r* ≃ 0.7 for all SNR >15 dB (Figure 4D). For SNR < 10 dB, the *r* value dropped to a correlation of *r*≃ 0.5. Despite the lower values for *r*, the mean curvature errors were reasonable. Relative to a true mean curvature of *κ* = 0.226 *μm*^−1^, at low SNR (< 10 dB) the curvature error is 0.04 *μm*^−1^ and decreases by two fold (0.02 *μm*^−1^) for higher SNR values (> 20 dB) (Figure 4E).

#### Variable Pixel Size

In addition to noise, the digitization of an imaging system may also limit accuracy and precision. Digital images are lossy: taking an image involves representing a continuous structure as a set of discrete pixels. Consequently, more finely discretized images may be less prone to error. We varied the pixel size from 0.1 to 0.4 *μ*m/pixel in a set of images with a fixed SNR of 20 dB (Figure 5). Fitting and error measurements were performed as described above. Over a broad range of pixel sizes, which spans most commonly used microscope cameras, we observed consistently strong fitting performance. E.g., Figure 5C and D show a plot of Pearson’s *r* against pixel size. The *r* value is very similar throughout, with decreases in performance occuring only when the pixel size was equal to or larger than the *σ* of the PSF. We conclude that pixel size is not a significant driver of fitting performance, at least at high SNR.

**Figure 5.**
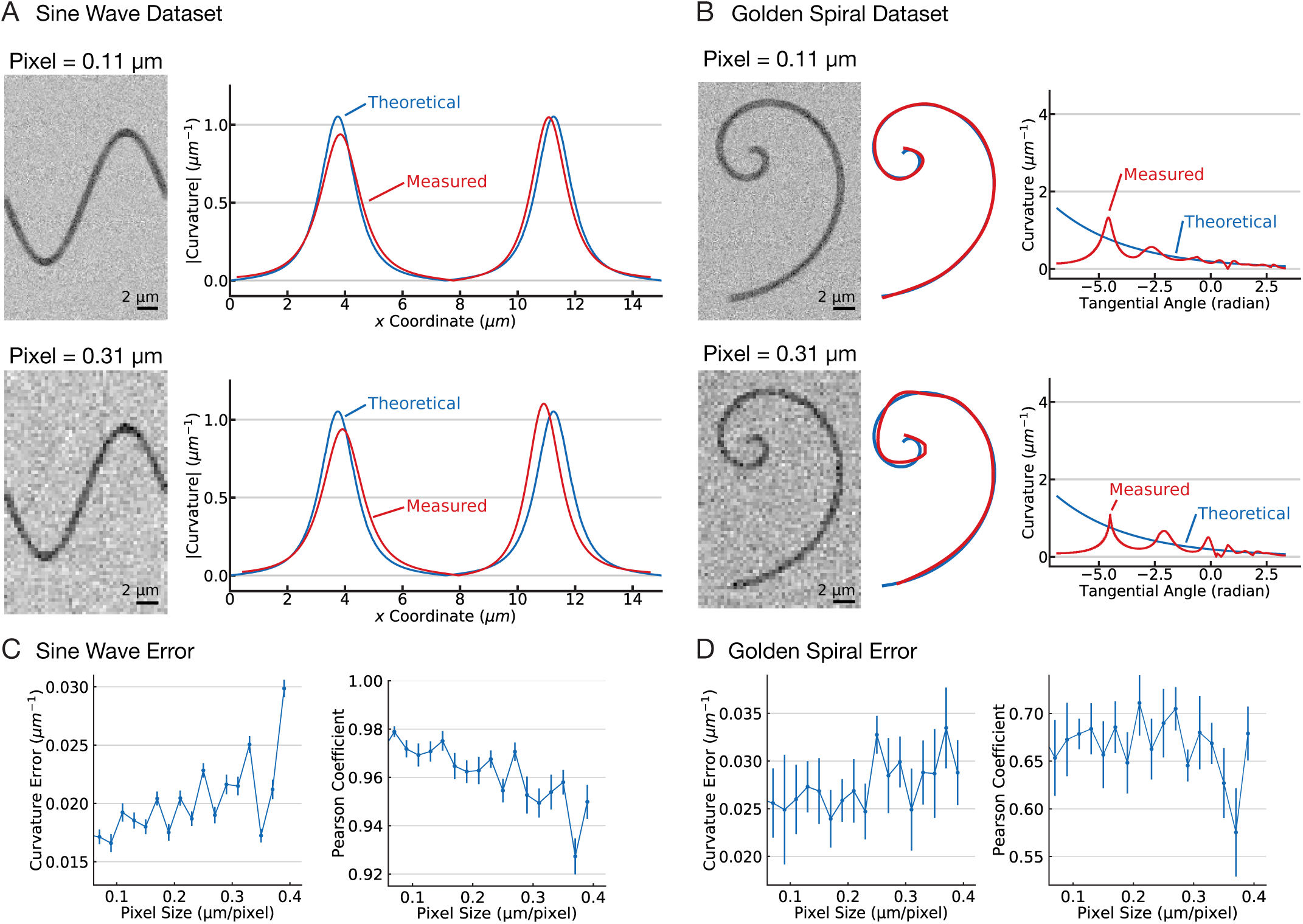
Fitting errors are reduced with smaller pixel sizes. **(A)** (*Left*) Synthetic images of sine waves at two pixel sizes (*labeled*). (*Right*) Plot of sine wave curvature against the *x* coordinate of the curve. The theoretical values are shown in blue and the measured values from a single measurent are shown in red. **(B)** (*Left*) Synthetic images of golden spirals at two pixel sizes (*labeled*). (*Right*) Plot of golden spiral curvature against the tangent angle of the curve (equivalent to a curve length coordinate). The theoretical values are shown in blue and the measured values from a single measurent are shown in red. **(C)** Error in *n* = 30 independent measurements of sine waves. (*Left*) Plot of the error in the mean curvature as a function of pixel size. Error bars are the SEM. (*Right*) Plot of the Pearson correlation coefficient as a function of pixel size. Error bars are the SEM. **(D)** Error in *n* = 30 independent measurements of golden spirals. (*Left*) Plot of the error in the mean curvature as a function of pixel size. Error bars are the SEM. (*Right*) Plot of the Pearson correlation coefficient as a function of pixel size. Error bars are the SEM.

#### Variable Initialization Curve

As mentioned above, our measurements of the sine wave data set used *n* = 30 independent initialization curves. These curves were meant to replicate real-world measurement conditions, in which different users will click out different initialization curves. The results of these 30 measurements were not identical (hence the error bars on, e.g., Pearson’s *r* in Figures 4 and 5). The non-identical results indicate that the “fitting landscape” for our algorithm does not have a single global minimum but rather many, closely-related local minima. This observation raises the possibility that a problematic initialization curve could hop out of these local minima entirely. In order to test Kappa against highly-variable initialization curves, we used sine waves and golden spirals with a SNR of 20 dB, pixel size of 0.16 *μ*m/pixel, and *σ* = 300 nm. We then added Gaussian noise to the *initial position* of our control points. The noise varied from 0.1 to 1 *μ*m for each control point of the initialization curves. Figures 6C and 6D show that fitting performance indeed breaks down when the initialization curves deviate significantly from the data, although the reduction in performance is smaller than we observed previously at low SNR. We conclude that, as long as Kappa can capture the correct pixels composing the curved object, our fitting procedure is able to find one of the local minima in the fitting landscape.

**Figure 6.**
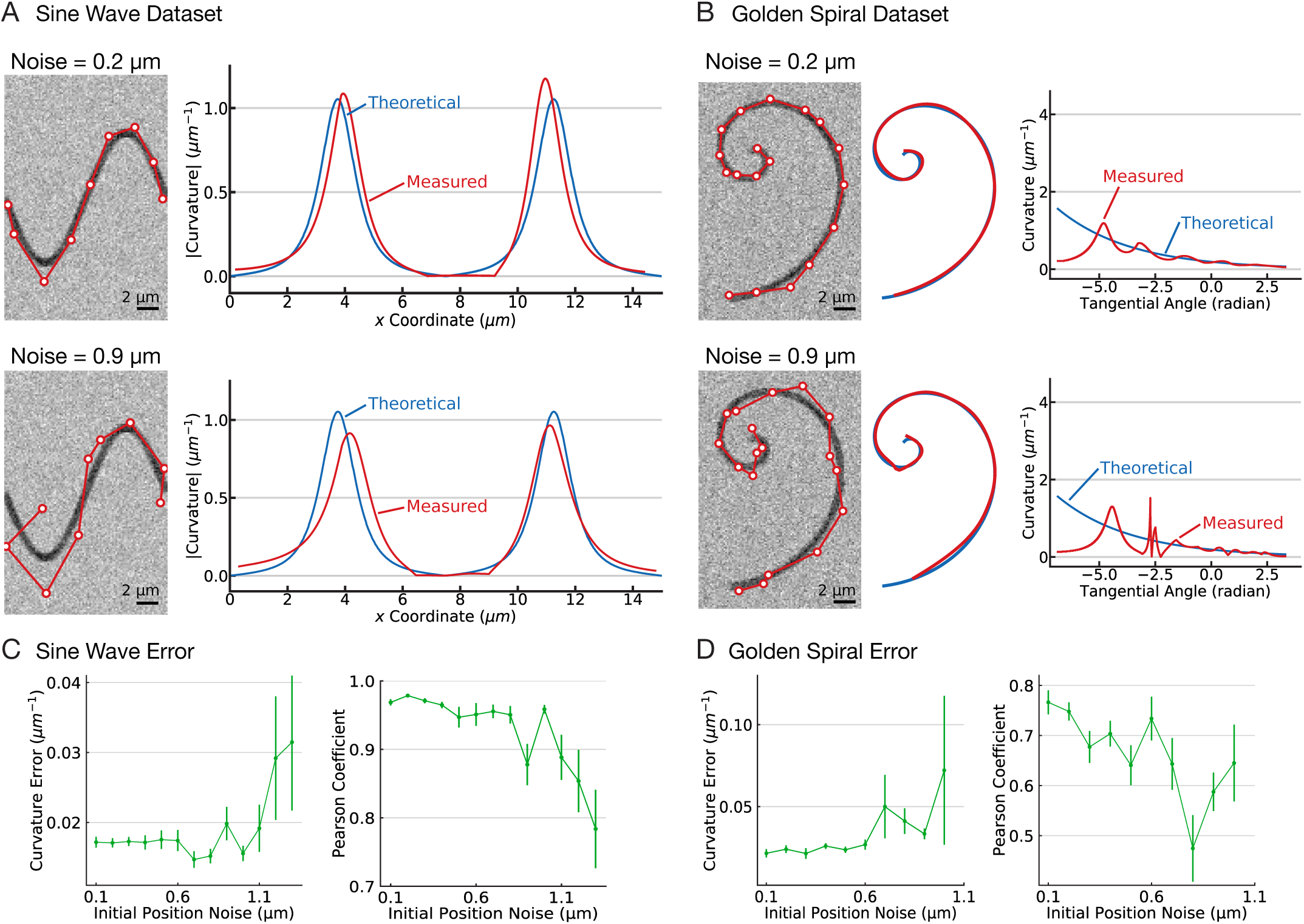
Fitting errors are insensitive to small differences in the initialization curves. **(A)** (*Left*) Synthetic images of sine waves showing two different initialization curves (*labeled*). (*Right*) Plot of sine wave curvature against the *x* coordinate of the curve. The theoretical values are shown in blue and the measured values from a single measurent are shown in red. **(B)** (*Left*) Synthetic images of golden spirals showing two different initialization curves (*labeled*). (*Right*) Plot of golden spiral curvature against the tangent angle of the curve (equivalent to a curve length coordinate). The theoretical values are shown in blue and the measured values from a single measurent are shown in red. **(C)** Error in *n* = 30 independent measurements of sine waves. (*Left*) Plot of the error in the mean curvature as a function of initial position noise. Error bars are the SEM. (*Right*) Plot of the Pearson correlation coefficient as a function of initial position noise. Error bars are the SEM. **(D)** Error in *n* = 30 independent measurements of golden spirals. (*Left*) Plot of the error in the mean curvature as a function of initial position noise. Error bars are the SEM. (*Right*) Plot of the Pearson correlation coefficient as a function of initial position noise. Error bars are the SEM.

#### Variable Point Spread Function

Finally, we tested how Kappa performs when the size of the PSF is varied. In the datasets described above, we convolved our images with a 2D Gaussian of *σ* = 300 nm, corresponding to *λ* = 530 nm light. These images were intended to replicate conventional fluorescence microscopy on green fluorescent proteins. In comparison, super resolution microscopy techniques can reduce the size of the PSF, either using advanced laser techniques (e.g., stimulated emission-depletion microscopy, where *σ* ≃ 100 nm) or using optical switching and fitting of single molecules (e.g., stochastic optical reconstruction microscopy, where *σ* is the error in the 2D Gaussian fit on each single molecule and is 20-50 nm). In order to test Kappa on “super resolution” images, we reduced the PSF size from 300 nm to 10 nm. This reduction produced the single largest improvement in performance (Figure 7). E.g., with *σ* values similar to STORM images, Kappa was nearly perfect, and the *x*-offset of the sine wave fits nearly disappeared. Similarly, the golden spiral fits more rapidly converged to smooth curves. We conclude that Kappa will perform optimally on super resolution images.

**Figure 7.**
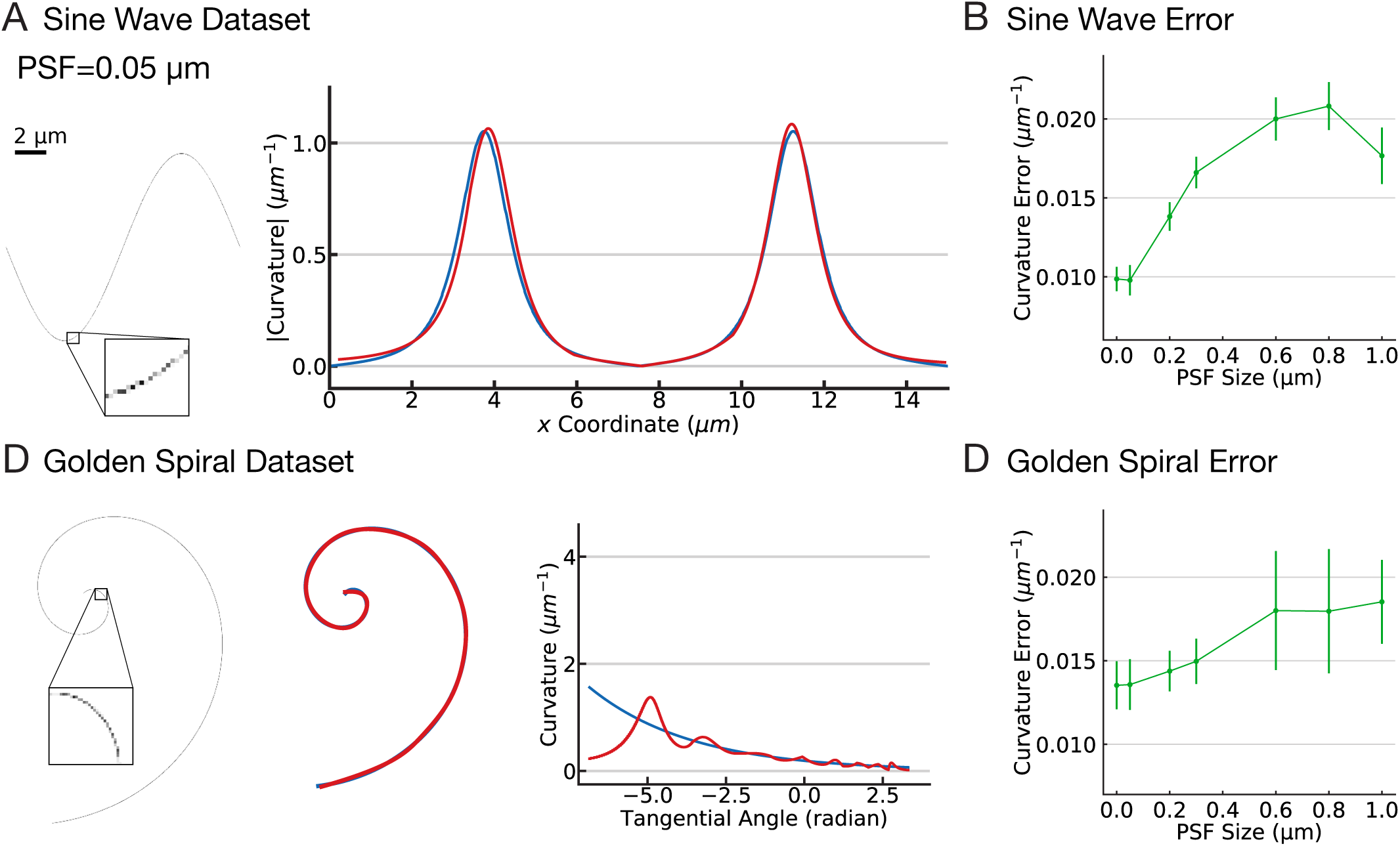
Fitting errors are significantly reduced with smaller PSF. **(A)** (*Left*) Synthetic image of a sine wave with a PSF equivalent to super-resolution microscopy data (*labeled*). (*Right*) Plot of sine wave curvature against the *x* coordinate of the curve. The theoretical values are shown in blue and the measured values from a single measurent are shown in red. **(B)** (*Left*) Synthetic images of a golden spiral with a PSF equivalent to super-resolution microscopy data (*labeled*). (*Right*) Plot of golden spiral curvature against the tangent angle of the curve (equivalent to a curve length coordinate). The theoretical values are shown in blue and the measured values from a single measurent are shown in red. **(C)** Error in *n* = 30 independent measurements of sine waves. Plot of the error in the mean curvature as a function of PSF. Error bars are the SEM. **(D)** Error in *n* = 30 independent measurements of golden spirals. (*Left*) Plot of the error in the mean curvature as a function of PSF. Error bars are the SEM.

## Availability and Reproducibility

Kappa is freely available on GitHub under the opensource MIT license. Kappa is a Fiji (Schindelin et al., 2012) plugin and so can be easily installed trough an ImageJ update site (check the manual for instructions). Anyone is free to submit bug reports and also code modifications to report issues and/or add new features to Kappa.

The source code of the `pykappa` Python library as well as the Jupyter notebooks used to generate the results presented in the Validation section are available at https://github.com/brouhardlab/Kappa/tree/master/Analysis. From this source code, anyone can reproduce our results and it can also serve as a good starting point to develop your own workflow in order to fit and measure curvatures of any objects on your own images. LaTeX sources of this manuscript are also available in the same repository.

## Discussion

Kappa is an easy-to-use and powerful tool to measure the curvatures of objects in images. Because Kappa is open-source, any developer can modify it. Below, we discuss some ideas for how Kappa might be further developed.

Kappa is an ImageJ1 (Schneider et al., 2012) plugin. The next generation of ImageJ, ImageJ2 (Rueden et al., 2017), was recently released and its usage is growing within the scientific imaging community. Thus, a natural evolution of Kappa would be switching its code base to ImageJ2, including porting the various algorithms presented in this paper as ImageJ Ops (Rueden et al., 2017). The switch would be an opportunity to work on computational speed, since Kappa slows down when the number of curves and control points becomes large (> 40). Another benefit of the switch would be an improvement in Kappa’s scripting capability. It is currently possible to script Kappa, but ImageJ1 requires that its user interface is loaded in memory even when executed from a script. ImageJ2 would not require this, because ImageJ Ops imposes a strict separation between user interface and logic.

Additional B-spline fitting algorithms could be implemented, such as the tangent distance minimization (Blake et al., 2012). Our two fitting algorithms struggled with the golden spiral (e.g., Fig. 6C) but other algorithms might perform better. Indeed, the best fitting algorithm may depend on the images being analyzed. We encourage all users to develop artificial data of their curved objects based on their own pixel size, SNR, and PSF, and then to benchmark the fitting algorithms accordingly.

The future of digital image analysis may (or may not) lie with machine learning, neural networks, and deep AI. Until such time as Skynet launches, we hope this program will prove useful.

## Acknowledgments

We thank K. Lu for many things: for developing early versions of code, for rickrolling us in the initial users manual, and for his beautiful charactature illustrations of folks in the lab. We thank C. Edgrington and S. Cruz Tetlalmatzi for their helpful comments on the manuscript. We thank E. Nazarova and J. Vogel for images of budding yeast. We thank M. Ahmadi and R. Roy for images of *C. elegans*. We thank S. Bechstedt for images of curved microtubules. G. Brouhard acknowledges support from Natural Sciences and Engineering Research Council of Canada (#RGPIN-2014-03791), the Canadian Institutes of Health Research (MOP-137055 and PJT-148702), and McGill University.

## Appendix

**Figure S1.**
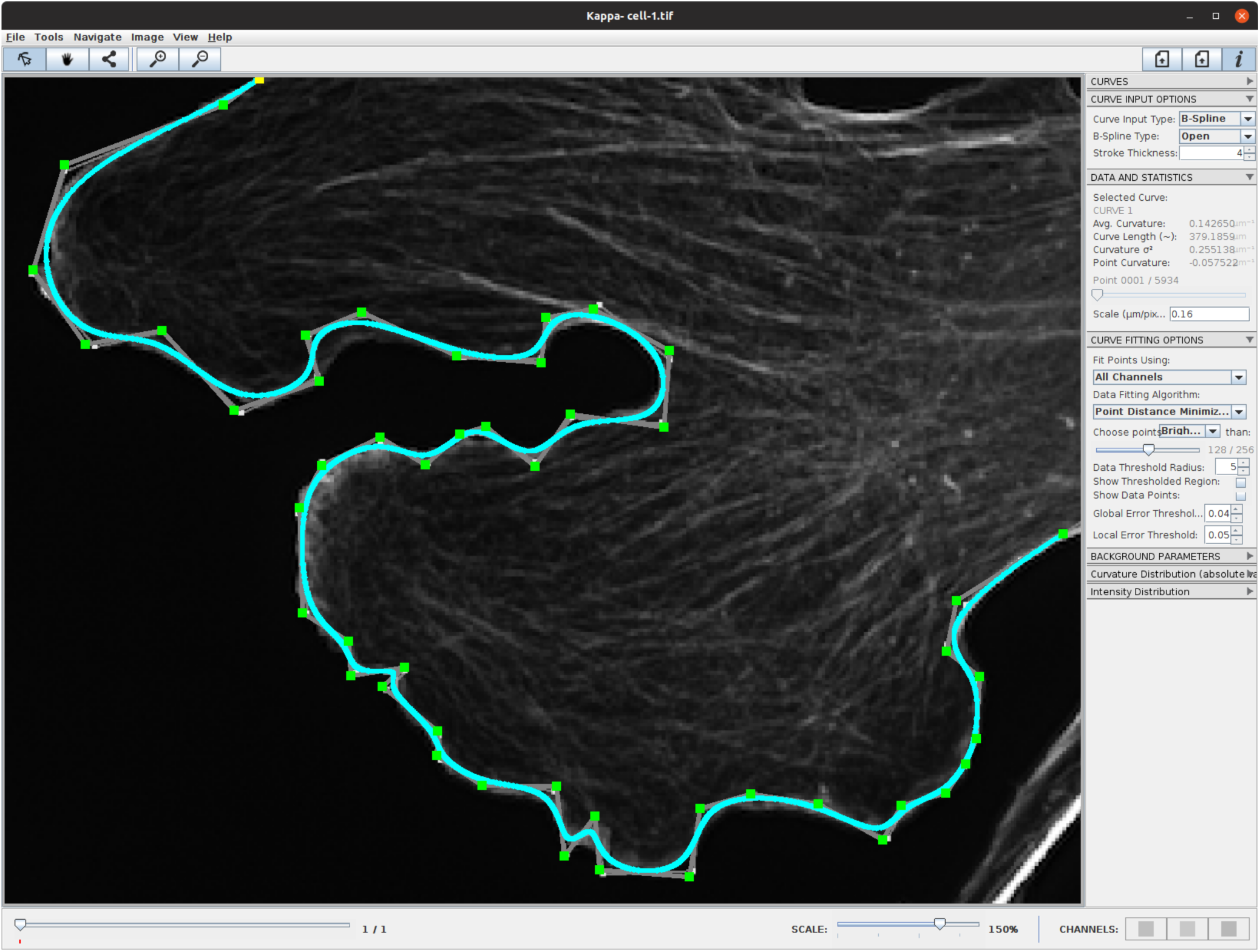
The user interface of Kappa. The top bar allows the user to draw, fit and modify curve using control points. The right panel contains multiples widget guiding the user through different steps; from drawing the initialization curves to the visualization of the curve’s curvature and associated pixel intensities. The bottom bar permits zoom-in and out as well as changing the current frame when working with a multistack image.

## References

Bah, T. (2011). Inkscape: guide to a vector drawing program. Vol. 559. Prentice Hall Upper Saddle River, NJ, USA.

Bechstedt, S., K. Lu, and G. Brouhard (Oct. 2014). “Doublecortin Recognizes the Longitudinal Curvature of the Microtubule End and Lattice”. Current Biology 24.20, pp. 2366–2375.

Berridge, M. J. (Nov. 2002). “The endoplasmic reticulum: a multifunctional signaling organelle”. Vol. 32. 5–6. Elsevier BV, pp. 235–249.

Bézier, P. E. (Feb. 1968). How Renault Uses Numerical Control for Car Body Design and Tooling. Tech. rep.

Bicek, A. D. et al. (2007). “Analysis of Microtubule Curvature”. Methods in Cell Biology. Elsevier, pp. 237–268.

Blake, A. and M. Isard (2012). Active contours: the application of techniques from graphics, vision, control theory and statistics to visual tracking of shapes in motion. Springer Science & Business Media.

Böhm, W. (1981). “Generating the Bézier points of Bspline curves and surfaces”. Computer-Aided Design 13.6, pp. 365–366.

Boor, C. de (1978). A Practical Guide to Splines. Springer New York.

Brangwynne, C. P. et al. (2006). “Microtubules can bear enhanced compressive loads in living cells because of lateral reinforcement”. The Journal of Cell Biology 173.5, pp. 733–741.

Bridges, A. A. et al. (Apr. 2016). “Micron-scale plasma membrane curvature is recognized by the septin cytoskeleton”. The Journal of Cell Biology 213.1, pp. 23–32.

Davis, P. J. (1975). Interpolation and approximation. Courier Corporation.

Eilers, P. H. and B. D. Marx (1996). “Flexible smoothing with B-splines and penalties”. Statistical science, pp. 89–102.

Field, C. M. (May 1996). “A purified Drosophila septin complex forms filaments and exhibits GTPase activity”. The Journal of Cell Biology 133.3, pp. 605–616.

Grossman, M. (Feb. 1971). “Parametric curve fitting”. The Computer Journal 14.2, pp. 169–172.

Hoschek, J. (July 1987). “Approximate conversion of spline curves”. Computer Aided Geometric Design 4.1-2, pp. 59–66.

Hoschek, J. (June 1988). “Intrinsic parametrization for approximation”. Computer Aided Geometric Design 5.1, pp. 27–31.

Hoschek, J., D. Lasser, and L. L. Schumaker (1993). Fundamentals of computer aided geometric design. AK Peters, Ltd.

Marsh, M. (July 1999). “The Structural Era of Endocytosis”. Science 285.5425, pp. 215–220.

McMahon, H. T. and E. Boucrot (July 2011). “Molecular mechanism and physiological functions of clathrin-mediated endocytosis”. Nature Reviews Molecular Cell Biology 12.8, pp. 517–533.

Pan, F., R. L. Malmberg, and M. Momany (2007). “Analysis of septins across kingdoms reveals orthology and new motifs”. BMC Evolutionary Biology 7.1, p. 103.

Plass, M. and M. Stone (1983). “Curve-fitting with piece-wise parametric cubics”. Proceedings of the 10th annual conference on Computer graphics and interactive techniques - SIGGRAPH ‘83. ACM Press.

Pressley, A. (2010). Elementary Differential Geometry. Springer London.

Risca, V. I. et al. (Jan. 2012). “Actin filament curvature biases branching direction”. Proceedings of the National Academy of Sciences 109.8, pp. 2913–2918.

Rogers, D. F. (1977). B-spline curves and surfaces for ship hull definition.

Roostalu, J., N. I. Cade, and T. Surrey (Sept. 2015). “Complementary activities of TPX2 and chTOG constitute an efficient importin-regulated microtubule nucleation module”. Nature Cell Biology 17.11, pp. 1422–1434.

Rueden, C. T. et al. (Nov. 2017). “ImageJ2: ImageJ for the next generation of scientific image data”. BMC Bioinformatics 18.1.

Samsonov, A. et al. (Dec. 2004). “Tau interaction with microtubules in vivo”. Journal of Cell Science 117.25, pp. 6129–6141.

Schindelin, J. et al. (June 2012). “Fiji: an open-source platform for biological-image analysis”. Nature Methods 9.7, pp. 676–682.

Schneider, C. A., W. S. Rasband, and K. W. Eliceiri (June 2012). “NIH Image to ImageJ: 25 years of image analysis”. Nature Methods 9.7, pp. 671–675.

Sederberg, T. W. (2012). Computer aided geometric design. BYU Faculty Publications.

Shibata, Y., T. Shemesh, et al. (Nov. 2010). “Mechanisms Determining the Morphology of the Peripheral ER”. Cell 143.5, pp. 774–788.

Shibata, Y., G. K. Voeltz, and T. A. Rapoport (Aug. 2006). “Rough Sheets and Smooth Tubules”. Cell 126.3, pp. 435–439.

Thomas, K. et al. (2016). “Jupyter Notebooks: a publishing format for reproducible computational workflows”. Stand Alone 0.Positioning and Power in Academic Publishing: Players, Agents and Agendas, pp. 87–90.

Wang, W., H. Pottmann, and Y. Liu (2006). “Fitting B-spline curves to point clouds by curvature-based squared distance minimization”. ACM Transactions on Graphics (ToG) 25.2, pp. 214–238.

Worring, M. and A. Smeulders (Nov. 1993). “Digital Curvature Estimation”. CVGIP: Image Understanding 58.3, pp. 366–382.

Yang, H., W. Wang, and J. Sun (June 2004). “Control point adjustment for B-spline curve approximation”. Computer-Aided Design 36.7, pp. 639–652.

